# DINC-COVID: A webserver for ensemble docking with flexible SARS-CoV-2 proteins

**DOI:** 10.1101/2021.01.21.427315

**Authors:** Sarah Hall-Swan, Dinler A. Antunes, Didier Devaurs, Mauricio M. Rigo, Lydia E. Kavraki, Geancarlo Zanatta

## Abstract

**Motivation:** Recent efforts to computationally identify inhibitors for SARS-CoV-2 proteins have largely ignored the issue of receptor flexibility. We have implemented a computational tool for ensemble docking with the SARS-CoV-2 proteins, including the main protease (Mpro), papain-like protease (PLpro) and RNA-dependent RNA polymerase (RdRp).

**Results:** Ensembles of other SARS-CoV-2 proteins are being prepared and made available through a user-friendly docking interface. Plausible binding modes between conformations of a selected ensemble and an uploaded ligand are generated by DINC, our parallelized meta-docking tool. Binding modes are scored with three scoring functions, and account for the flexibility of both the ligand and receptor. Additional details on our methods are provided in the supplementary material.

**Availability:** dinc-covid.kavrakilab.org

**Supplementary information:** Details on methods for ensemble generation and docking are provided as supplementary data online.

## 1 Introduction

The novel coronavirus SARS-CoV-2, which causes the respiratory disease COVID-19, went from an outbreak to a pandemic in just a few months (Rabaan *et al*., 2020). In response, there have been unprecedented global efforts to develop effective treatments. Among pharmacological targets, proteins involved in the viral replication have been used in several computational studies focused on drug design, drug repurposing and virtual screening. Unfortunately, proposed SARS-CoV-2 inhibitors have not yet impacted the course of the COVID-19 pandemic.

Although an impressive number of SARS-CoV-2 proteins have been solved in a short period of time, computational efforts targeting these proteins have been limited to the use of a single experimental structure. Such approach is expected to correctly reproduce similar experimental structural data, but tends to fail in exploring the binding of ligands with diverse chemical features (i.e., larger ligands or alternative binding modes) as it do not account for the inherent flexibility of proteins in solution (Kneller et al., 2020).

Here we present DINC-COVID, a webserver that accounts for the flexibility of both ligands and proteins in determining binding modes of small molecules and peptides interacting with SARS-CoV-2 proteins (Fig. 1). DINC-COVID is based on DINC (Devaurs et al., 2019), a parallelized meta-docking approach that was shown to outperform conventional docking tools on several challenging datasets, especially for large and flexible ligands (e.g., peptidomimetics and peptides).

**Fig. 1.**
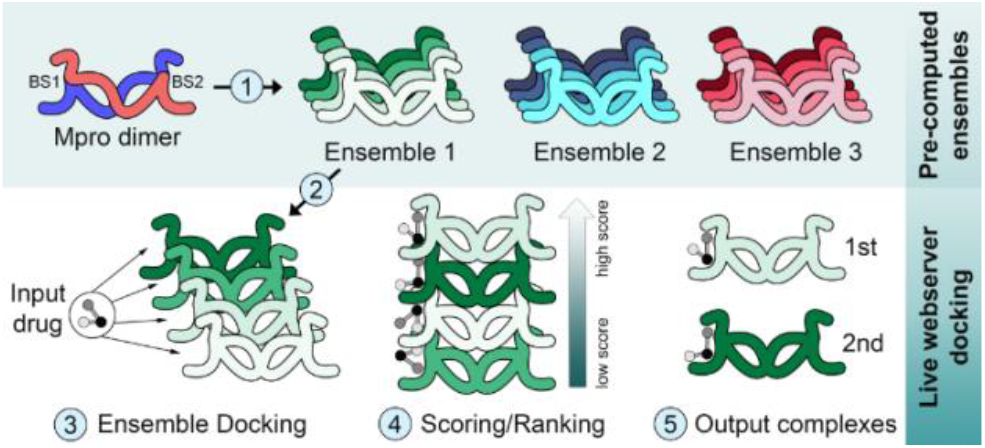
Ensemble docking with DINC-COVID. The top left figure is a schematic representation of the SARS-CoV-2 main protease (MPro) dimeric structure, with one protomer depicted in blue, and another in red. 1. Ensemble generation: Three ensembles of the Mpro dimer were pre-computed (see Supplementary Methods) and made available through the DINC-COVID webserver. Different shades represent different conformations within each ensemble. 2. Input selection: The user can select one of the available ensembles, and upload a ligand of interest (e.g., drug or peptide). 3. Ensemble docking: The parallelized meta-docking approach DINC is used to sample alternative binding modes using all the receptor conformations in the ensemble. 4. Scoring and Ranking: All generated binding modes are rescored and ranked using three scoring functions (i.e., Vina, Vinardo and AutoDock4). 5. Output complexes: The top scoring complexes are returned to the user, reflecting the flexibility of both the ligand and receptor. BS, binding site.

DINC-COVID users have the option to choose among three distinct ensembles for each SARS-CoV-2 target protein, including data from experimental structures (crystal) or from molecular dynamics with distinct force fields (charmm or gromos). Generating representative ensembles of protein conformations is a challenging and computationally intensive task, thus each ensemble was tailored through robust data analysis and sampling methodology, reflecting different degrees of flexibility and features of the targeted binding site. By making DINC-COVID available as a webserver with pre-computed ensembles, we aim to facilitate and speed-up the use of ensemble docking in the search of efficient SARS-CoV-2 inhibitors (Fig. 1).

## 2 DINC-COVID: A User-friendly Webserver

### 2.1 User interface

An intuitive interface allows users to upload a ligand and select the receptor ensemble (Fig. S1). Three ensembles for each target site are currently available for docking (see details in Supplementary Methods). Optional ligand preparation (i.e., adding hydrogens, charges, etc) is also available. Finally, all generated binding modes are rescored and ranked using three scoring functions (i.e.,Vina, Vinardo and AutoDock4), outputting a user-defined number of top-scoring conformations. The “Results” page provides the top-scoring binding modes, including both the ligand and the receptor (Fig. S1). Users can view the conformations in the embedded visualization tool, and download the results for offline analysis. Downloadable files include top-scoring complexes in pdb format, and a text file listing selected conformations and binding energies.

### 2.2 Validation

At the time of our study, 116 out of 156 Mpro crystal structures were complexed with ligands. However, in 80 of these structures the ligand was covalently bound, leading to high energy binding modes that cannot be reproduced. The remaining 36 Mpro structures were selected for self-docking and ensemble docking experiments (Table S5). When self-docking, DINC-COVID was able to produce near-native binding modes for all complexes, as determined by the root-mean square deviation (RMSD) between the binding modes and the ligand crystal structure (e.g., mean heavy-atom RMSD of only 1.4 Å). Note that the goal of ensemble-docking is not to reproduce rigid crystal structures, but to find alternative lower energy binding modes. Therefore, self-docking is not the best validation approach, since by design the top ranking conformation will often not have the lowest RMSD to the crystal. In fact, different receptor conformations contribute to the lower energy binding modes in different ensemble docking experiments (Fig. S7). These results highlight the potential of DINC-COVID to identify completely novel binding modes, and to reveal yet unknown candidate inhibitors. In this context, we tested our predictive power against a set of experimentally characterized Mpro inhibitors (Table S6) and antiviral peptidomimetics (Table S7). As noted, the binding of these peptidomimetics requires malleability of Mpro catalytic site (Kneller *et al*., 2020). In both cases, we obtained good correlation with experimental data (r = 0.88 and r = 0.86, respectively), and outperformed other existing single target docking solutions, especially when considering peptidomimetics.

## 3 Discussion

Our webserver offers a ready-to-use solution for researchers to account for protein flexibility while testing compounds against SARS-CoV-2 proteins. It allows users to run ensemble docking experiments, without the additional burden of time-consuming simulations required for ensemble generation and docking preparation. For instance, we have carefully tuned the protonation state of His, Glu and Asp residues for each protein conformation. Altogether, these features make our server of interest even for advanced users, adding robustness to docking analysis. Currently, DINC-COVID offers ensemble docking for the catalytic binding site of SARS-CoV-2 Main protease (Mpro), Papain-like protease (PLpro) and RNA-dependent RNA polymerase (RdRp) (Zhu *et al*., 2020; Aftab *et al*., 2020; Faheem *et al*., 2020), as well as the allosteric binding site of Mpro (Menéndez *et al*., 2020). Ensembles of other SARS-CoV-2 proteins are under process (e.g., spike protein) and will be added to the webserver, which also enables users to make requests. Finally, we are working on a version of DINC-COVID that can be deployed on local computational resources, to enable its use for virtual screening.

## Acknowledgements & Funding

We thank the Center for Research Computing (CRC) at Rice University for supporting our use of ORION VM Pool. Use of CRC resources is supported by the Data Analysis and Visualization Cyberinfrastructure funded by NSF (OCI-0959097) and by Rice University. We also thank the *Centro Nacional de Supercomputação* (CESUP/UFRGS, Brazil), whose resources were used to perform our MD simulations. This work was funded in part by the National Science Foundation IIBR:Informatics:RAPID program (2033262), the Cancer Prevention & Research Institute of Texas through a fellowship from the Gulf Cost Consortia on the Computational Cancer Biology Training Program (RP170593), and from the National Library of Medicine Training Program (T15LM007093-29). This work has also been supported in part by the National Council for Scientific and Technological Development (Brazil, 437373/2018-5), and by Rice University funds.

## References

Aftab, S.O. et al. (2020) Analysis of SARS-CoV-2 RNA-dependent RNA polymerase as a potential therapeutic drug target using a computational approach. J. Transl. Med.

Devaurs, D. et al. (2019) Using parallelized incremental meta-docking can solve the conformational sampling issue when docking large ligands to proteins. BMC Mol. Cell Biol.

Faheem et al. (2020) Druggable targets of SARS-CoV-2 and treatment opportunities for COVID-19. Bioorg. Chem.

Kneller, D.W. et al. (2020) Malleability of the SARS-CoV-2 3CL Mpro Active-Site Cavity Facilitates Binding of Clinical Antivirals. Structure.

Menéndez, C.A. et al. (2020) Molecular characterization of ebselen binding activity to SARS-CoV-2 main protease. Sci. Adv.

Rabaan, A.A. et al. (2020) SARS-CoV-2/COVID-19 and advances in developing potential therapeutics and vaccines to counter this emerging pandemic. Ann. Clin. Microbiol. Antimicrob.

Zhu, W. et al. (2020) RNA-Dependent RNA Polymerase as a Target for COVID-19 Drug Discovery. SLAS Discov.

