## Supplemental Information for "DINC-COVID: A webserver for ensemble docking with flexible SARS-CoV-2 proteins"

### DINC-COVID Algorithm

DINC-COVID uses parallel threads to execute independent sampling of the ligand with Autodock Vina(Trott and Olson, 2009). Then, output conformations are ranked using scoring functions from AutoDock Vina, AutoDock4 (Morris *et al.*, 2009), and Vinardo (Quiroga and Villarreal, 2016). The best binding modes for each scoring function are chosen, and the resulting complexes are returned (Fig. 1). More details are available at http://dinc-covid.kavrakilab.org/method/.

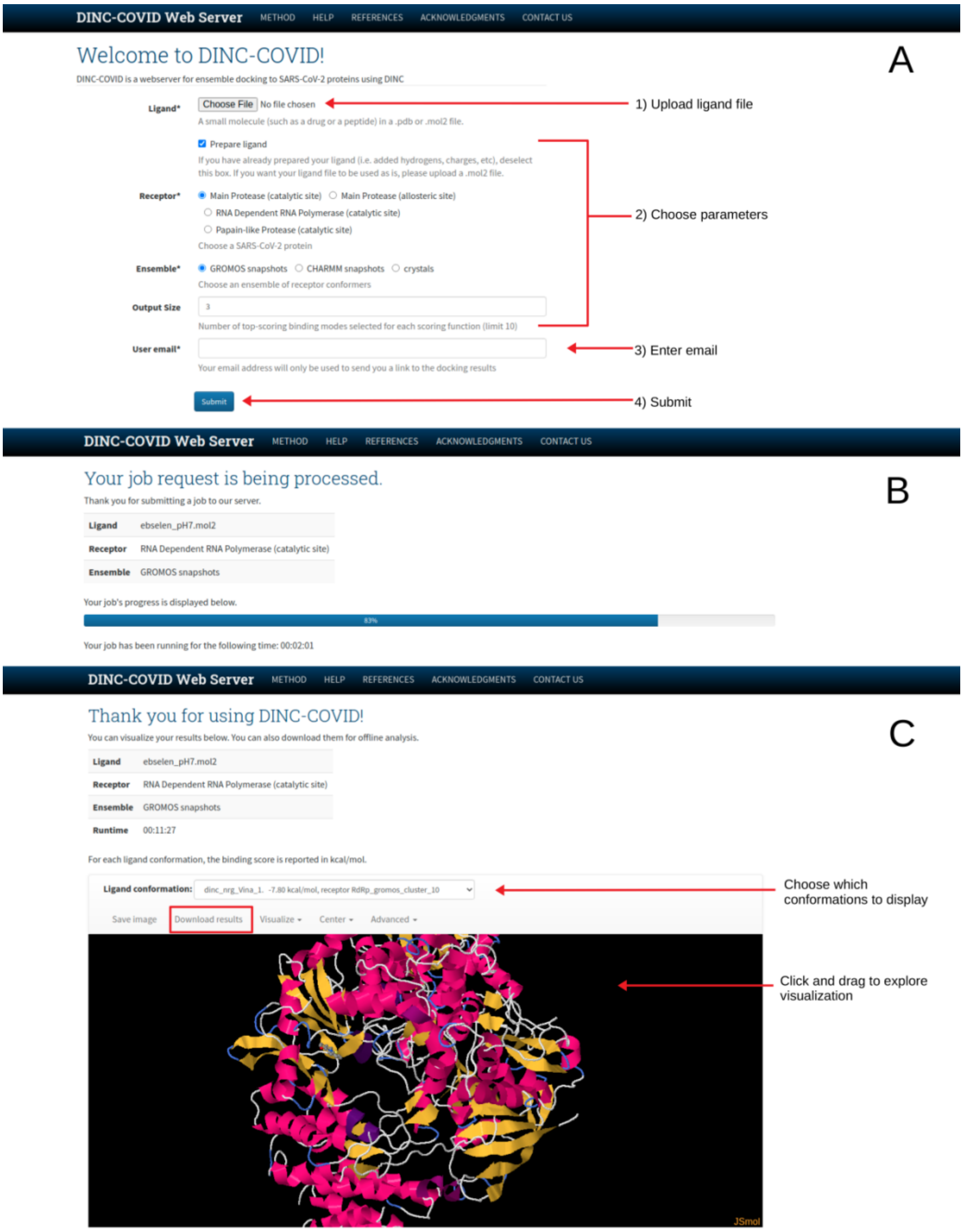


Supplementary Figure S1: Running ensemble docking with DINC-COVID. A. The home page of DINC-COVID allows the user to (1) upload ligand (i.e., pdb or mol2 file), (2) choose parameters (e.g., request preparation of the ligand and select receptor ensemble), (3) provide an e-mail address and (4) submit the job. B. A link is sent to the user by e-mail, to check the progress of the ongoing job. C. The same link allows checking the results in the DINC-COVID webpage. The user can browse through different binding modes, visualize the conformations, or download all results for further inspection.

**Ensemble Generation**

Four ensembles of SARS-CoV-2 proteins are currently available through the DINC-COVID webserver. For each target binding site, three ensembles were made available (Fig. 1 and S1A). The first one contains structures from experimental data. When the number of structures was large (i.e. by the time of this analysis, the main protease had 156 crystallographic structures available), representative conformations were sampled as described below. The other two ensembles are sets of conformations sampled from 2 µs of molecular dynamics (MD) simulations with the GROMACS 2019 package. When necessary, the biological assembly was produced with PDBePISA (Krissinel and Henrick, 2007). All structures were protonated at pH 7.0 using the PROPKA algorithm (Olsson *et al.*, 2011) at the PDB2PQR server (Dolinsky *et al.*, 2004). The choice of the force field has significant impacts on the results of MD simulations, since it encompasses a series of empirically-determined parameters (Lopes *et al.*, 2015; Villavicencio *et al.*, 2018). Therefore, we created separate ensembles with two force fields which use distinct representations for hydrogen atoms, to avoid limitations on conformational sampling due to force field bias. Two sets of MDs were executed using CHARMM (i.e., CHARMM36 (Best *et al.*, 2012)) and GROMOS (i.e., GROMOS53a6 (Oostenbrink *et al.*, 2004)) force fields, respectively (i.e., five runs of 200 ns for each force field). These ensembles reflect different scales of protein flexibility, from subtle side chain rearrangements (e.g., crystal ensemble), to larger backbone motions (e.g., gromos ensemble). To build the final ensembles, approximately 100,000 conformations (per set of MD) were extracted from trajectories using MDtraj (McGibbon *et al.*, 2015). Data reduction was performed using Principal Components (PCs) needed to explain ≈80% of variance. The free energy of these conformations was estimated with PyEMMA (Scherer *et al.*, 2015) and plotted over the first two PCs. The Elbow method was used to determine the ideal number of clusters (k) in each case. Finally, the K-means clustering algorithm from scikit-learn package (Pedregosa *et al.*, 2011) was employed to identify representative members for each cluster. Representative images of ensembles from crystallographic structures for the Main protease (Mpro), papain-like protease (PLpro) and the RNA-dependent RNA polymerase (RdRp) are shown in supplementary figure S2.

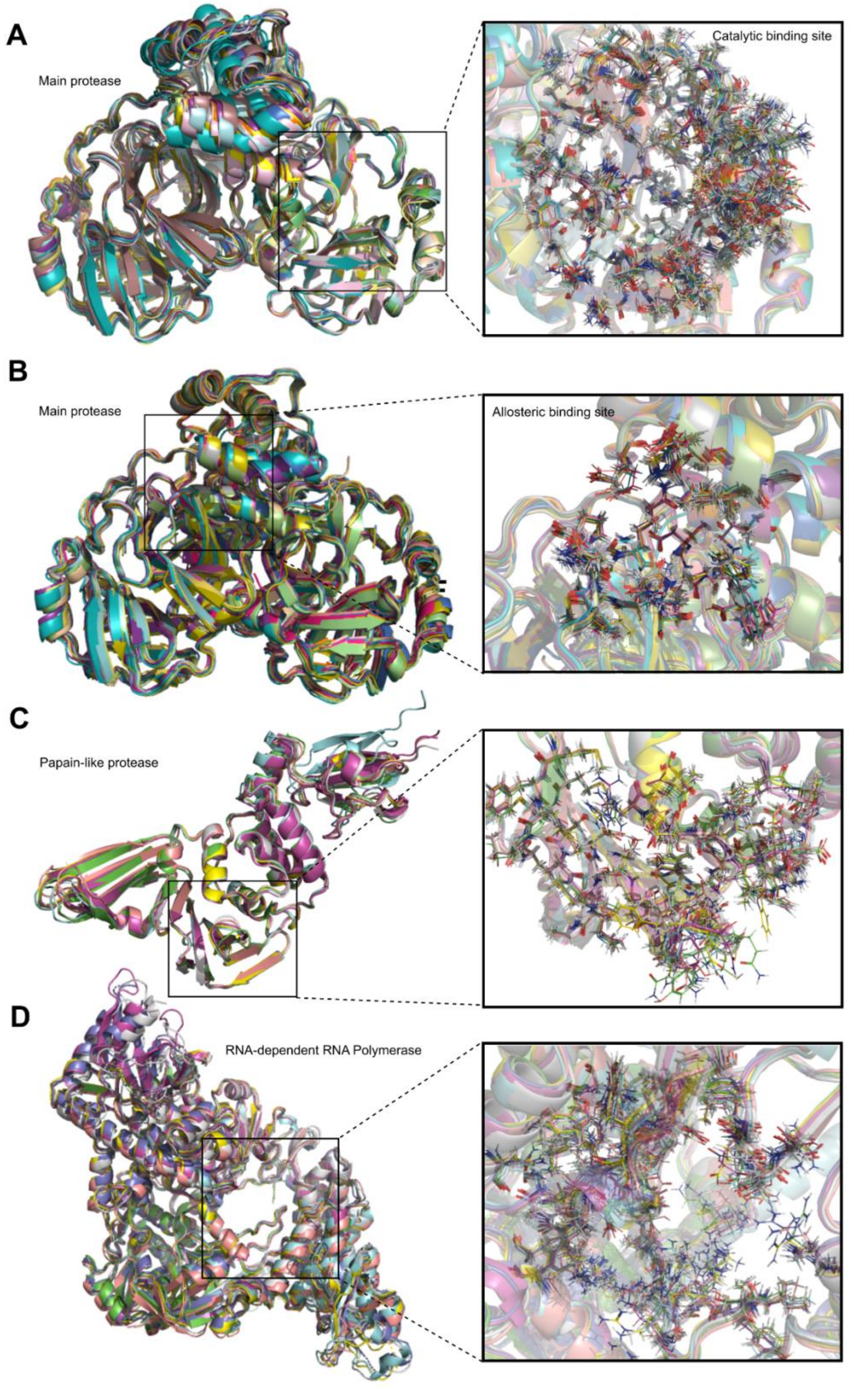


Supplementary Figure S2: Cartoon and sticks representation of the crystal ensembles for the four binding sites available at the DINC-COVID webserver. A. Catalytic binding site of Mpro. B. Allosteric binding site of Mpro. C. Catalytic binding site of PLpro. D. Catalytic binding site of RdRp. Even broader conformational flexibility is captured by the provided MD-derived ensembles.

**NSP5 - Main protease (Mpro, 3CLPro)**

For the ensemble of crystals, the dimeric state of 156 crystallographic structures was sampled based on conformational changes in the catalytic (supplementary figure S3A) or allosteric (supplementary figure S4A) binding pocket. In addition, two new ensembles were produced by sampling conformations from molecular dynamics trajectories. For MD simulations, the dimeric form of Mpro was built from structure 6LU7 and used as input.

<u>Catalytic binding site</u>: The selected Mpro crystal structures are shown in Table S1. The total number of structures in the ensemble of crystals, charmm simulations and gromos simulations are 25, 14 and 11, respectively (supplementary figure S3B-F). The scoring box center was set to -8.50, 13.40 and 67.80 (X, Y and Z, respectively) with dimensions 70x76x62 Å for crystal ensemble; -10.50, 13.13 and 66.80 (X, Y and Z, respectively) with dimensions 82x88x94 Å for charmm ensemble; and -10.50, 13.13 and 66.80 (X, Y and Z, respectively) with dimensions 84x92x100 Å for gromos ensemble. While the box used for scoring is the same for the entire ensemble, the box used for sampling was tuned to the binding site of each receptor conformation, based on the solvent accessible volume of the binding site.

<u>Allosteric binding site</u>: The amount of structures into the ensemble of crystals, charmm simulations and gromos simulations are 20, 10 and 09, respectively (supplementary figure S4B-F). The scoring box center was set to -39.23, 7.85 and 57.56 (X, Y and Z, respectively) with dimensions 82x78x82 Å for crystal ensemble; -36.23, 7.09 and 85.56 (X, Y and Z, respectively) with dimensions 82x76x82 Å for charmm ensemble; and -37.23, 6.09 and 58.56 (X, Y and Z, respectively) with dimensions 84x76x82 Å for gromos ensemble. The box used for sampling was tuned to the binding site of each receptor conformation, as described above.

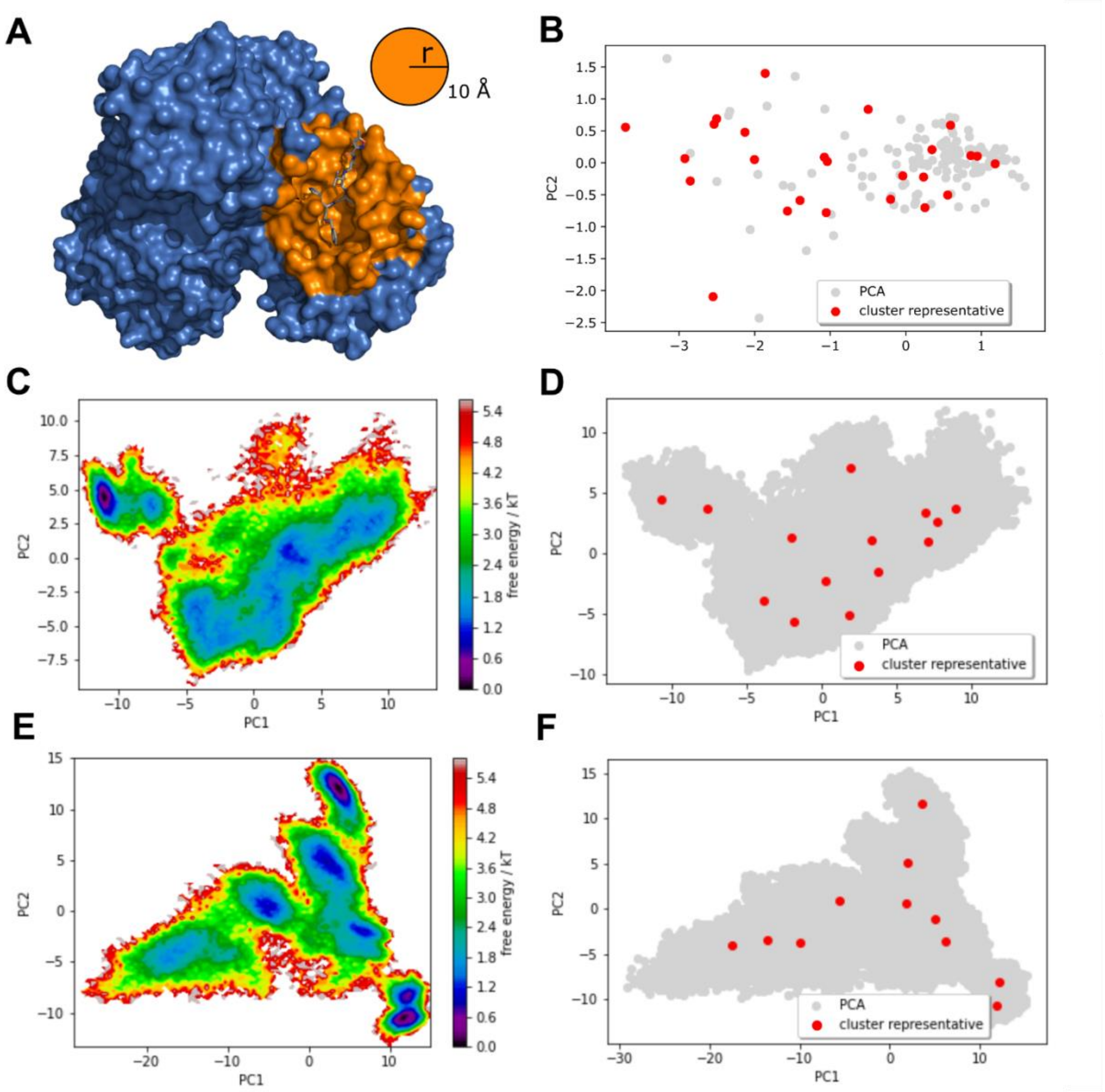


Supplementary Figure S3: Selection of representative conformations to create the Mpro Catalytic site ensembles. These ensembles take into account the flexibility of the catalytic binding site, using as reference all residues within 10 Å from the ligand N3 in the crystallographic structure 6LU7 (A). The conformational space for the unbound (i.e., apo) Mpro structure was sampled using data extracted from simulations using either available experimental structures (B), CHARMM36 force field (D) or GROMOS53a6 force field (F). The K-means clustering algorithm from scikit-learn package (Pedregosa *et al.*, 2011)[9] was employed to identify representative members for each cluster (red circles). The free energy of these conformations was estimated with PyEMMA (Scherer *et al.*, 2015) and plotted over the first two PCs (C, E) (i.e., darker colors indicate local minimas). The crystal structures ensemble represents the least amount of flexibility. When compared to the crystal ensemble, in terms of the range of Root Mean Square Deviation (RMSD) for the heavy atoms of the binding site 1 (BS1), the CHARMM ensemble presents intermediate flexibility (0.89-2.69 Å), while the GROMOS ensemble shows the largest variation (1.38-4.08 Å).

Supplementary table S1: Selected structures for the crystal-based Mpro catalytic site ensemble

| PDB ID | State | PubChem CID | Resolution (Å) | PDB ID | State | PubChem CID | Resolution (Å) |
|---|---|---|---|---|---|---|---|
| 5R84 | holo | 1072430 | 1.83 | 6M2N[a] | holo | 5281605 | 2.2 |
| 5RE9 | holo | 880785 | 1.72 | 6WNP | holo | 10324367 | 1.44 |
| 5RER | holo | 404914157 | 1.88 | 6XB0 | apo | - | 1.8 |
| 5RF5 | holo | 265635 | 1.74 | 6XBG | holo | - | 1.45 |
| 5RFA | holo | 1224835 | 1.52 | 6XCH | holo | 137348943 | 2.2 |
| 5RFH | holo | 60645778 | 1.58 | 6YVF | holo | 44137675 | 1.6 |
| 5RFJ | holo | 3511405 | 1.8 | 7BQY | holo | 146025593 | 1.7 |
| 5RFK | holo | 20754800 | 1.75 | 7BRO | apo | - | 2.2 |
| 5RGU | holo | 146037571 | 2.11 | 7BRP | holo | 10324368 | 1.8 |
| 5RH2 | holo | 89468951 | 1.83 | 7C8R | holo | 405559748 | 2.3 |
| 5RH6 | holo | 405243691 | 1.6 | 7C8T | holo | 11844232 | 2.05 |
| 5RH9 | holo | 405243697 | 1.91 | | | | |
| 6M0K | holo | 405243775 | 1.5 | | | | |

This subset of 25 conformations was selected with methods described in supplementary figure S3, from a total of 156 crystal structures including both apo (i.e., unbound) and holo (i.e., ligand-bound) Mpro conformations. [a] The crystallographic file 6M2N has two full dimers, with protomers showing alternative protonation states. Two of these states were selected and included in the final ensemble.

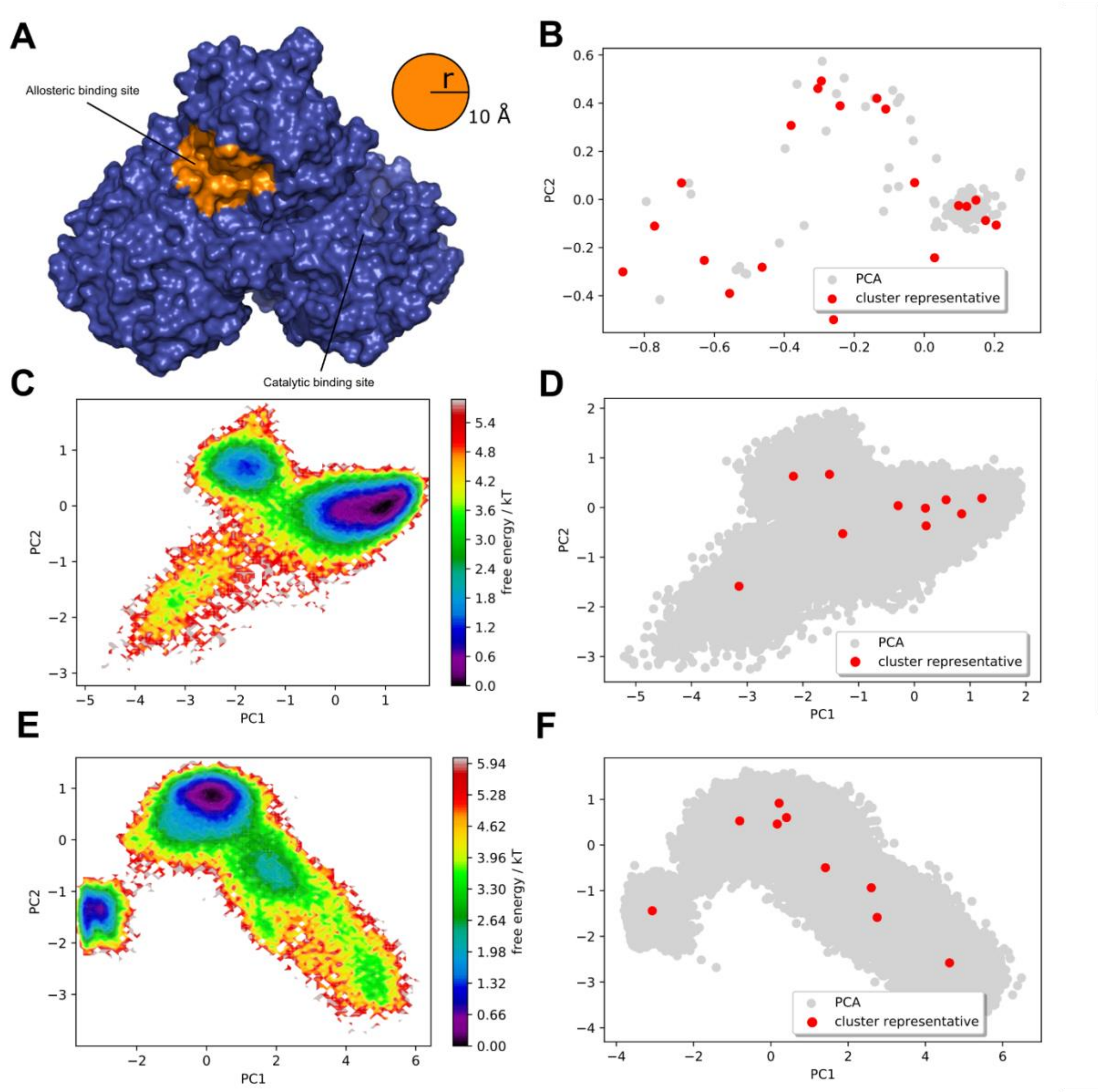


Supplementary Figure S4: Selection of representative conformations to create the Mpro allosteric binding site ensembles. These ensembles take into account the flexibility of the allosteric binding site, as indicated by (Menéndez *et al.*, 2020) (A). The conformational space for the unbound (i.e., apo) Mpro structure was sampled using data extracted from simulations using either available experimental structures (B), CHARMM36 force field (D) or GROMOS53a6 force field (F). The K-means clustering algorithm from scikit-learn package (Pedregosa *et al.*, 2011) was employed to identify representative members for each cluster (red circles). The free energy of these conformations was estimated with PyEMMA (Scherer *et al.*, 2015) and plotted over the first two PCs (C, E) (i.e., darker colors indicate local minimas).

Supplementary table S2: Selected structures for the crystal-based Mpro allosteric site ensemble

| PDB ID | State | PubChem CID | Resolution (Å) | PDB ID | State | PubChem CID | Resolution (Å) |
|---|---|---|---|---|---|---|---|
| 5r7y | holo | 118569 | 1.65 | 6wtt | holo | 137349627 | 2.15 |
| 5ref | holo | 2806372 | 1.61 | 6xb2 | holo | 16842 | 2.10 |
| 5reu | holo | 3803220 | 1.69 | 6xbh | holo | - | 1.60 |
| 5rfy | holo | 146018721 | 1.90 | 6xbi | holo | - | 1.70 |
| 5rgo | holo | 565340 | 1.74 | 6xch | holo | 137348943 | 2.20 |
| 5rgp | holo | 26865112 | 2.07 | 6xfn | holo | - | 1.70 |
| 5rh7 | holo | 146673002 | 1.71 | 6y2g | holo | 146018708 | 2.20 |
| 6m03 | apo | - | 2.00 | 6yvf | holo | 44137675 | 1.60 |
| 6m0k | holo | 146672237 | 1.50 | 7buy | holo | 14741611 | 1.60 |
| 6wtk | holo | 78225172 | 2.00 | 7c8u | holo | 137349627 | 2.35 |
| 5r7y | holo | 118569 | 1.65 | 6wtt | holo | 137349627 | 2.15 |
| 5ref | holo | 2806372 | 1.61 | 6xb2 | holo | 16842 | 2.10 |

This subset of 24 conformations was selected with methods described in supplementary figure S4, from a total of 156 crystal structures including both apo (i.e., unbound) and holo (i.e., ligand-bound) Mpro conformations. Note that all selected structures have the allosteric binding site in the apo state and all ligands listed are located in the catalytic binding site.

**NSP3 – Papain-like protease (PLpro)**

Three ensembles were built for the PLpro, based on conformational changes in the catalytic binding site. The target binding site was set based on structure 6WUU (supplementary figure S5A). For the ensemble of crystallographic structures, monomeric forms of PLpro were extracted from 12 crystals representing wild type structures (table S3). Crystals with double occupancy were split into two files to ensure each file would represent one distinct orientation. In addition, two new ensembles were produced by sampling conformations from molecular dynamics trajectories of PLpro in the APO state. In the case of MD, a monomeric unit from the crystallographic structure 6WUU was used as input, and approximately 100,000 conformations were analysed for each force field set of simulations. The amount of structures into the ensemble for crystals, charmm simulations and gromos simulations are 6, 20 and 19, respectively (supplementary figure S5B-F). The scoring box center was set to 46.9, 38.8 and 30.19 (X, Y and Z, respectively) with dimensions 78x80x82 Å for crystal ensemble; 48.1, 37.8 and 30.19 (X, Y and Z, respectively) with dimensions 78x80x82 Å for charmm ensemble; and 47.1, 37.8 and 31.19 (X, Y and Z, respectively) with dimensions 78x80x86 Å for gromos ensemble. The box used for sampling was tuned to the binding site of each receptor conformation, based on the solvent accessible volume of the binding site.

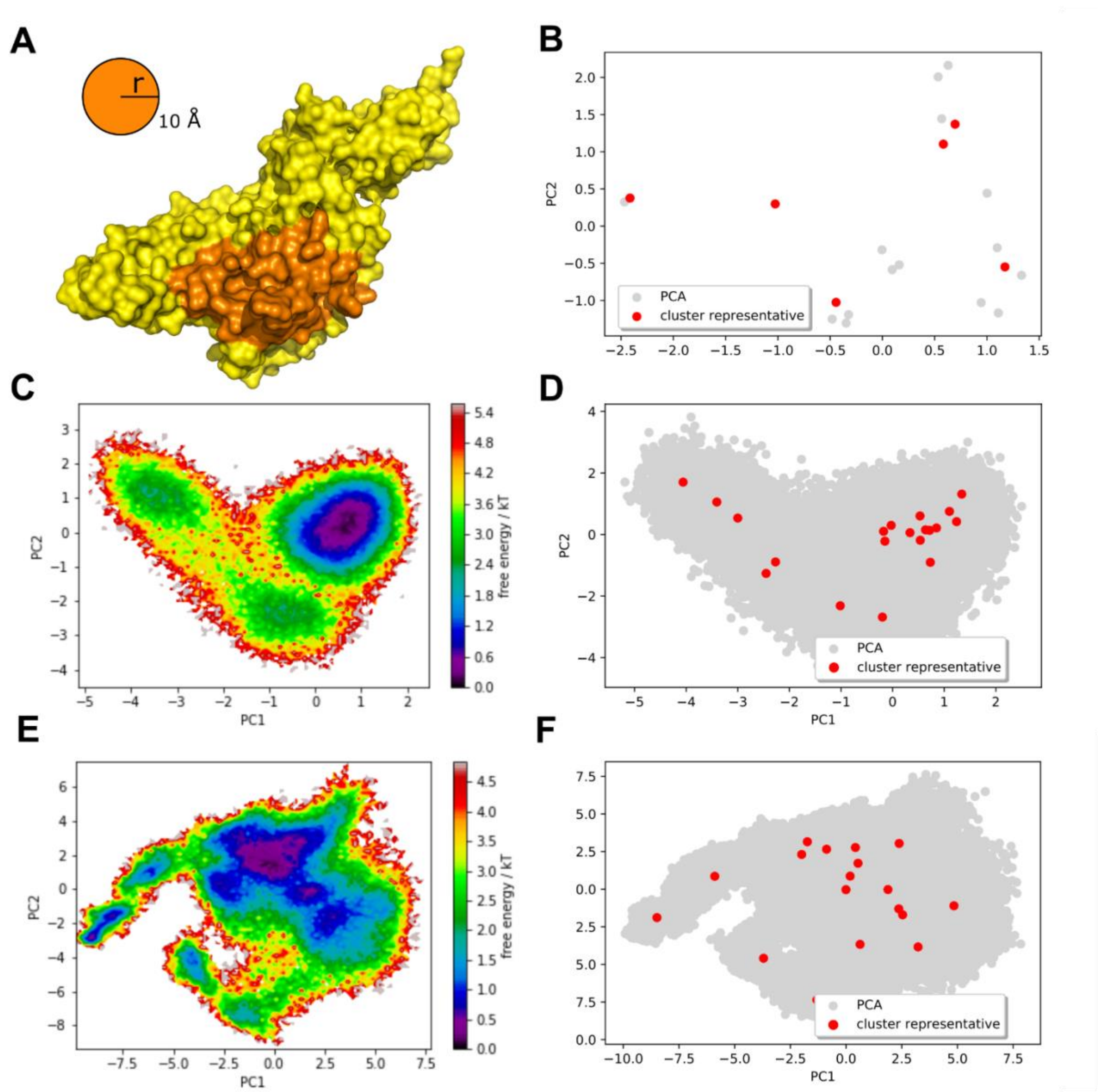


Supplementary Figure S5: Selection of representative conformations to create the PLpro ensembles. These ensembles take into account the flexibility of residues shown in orange (A). The conformational space for the unbound (i.e., apo) PLpro structure was sampled using data extracted from simulations using either available experimental structures (B), CHARMM36 force field (D) or GROMOS53a6 force field (F). The K-means clustering algorithm from scikit-learn package (Pedregosa *et al.*, 2011) was employed to identify representative members for each cluster (red circles). The free energy of these conformations was estimated with PyEMMA (Scherer *et al.*, 2015) and plotted over the first two PCs (C, E) (i.e., darker colors indicate local minimas).

Supplementary table S3: Selected structures for the crystal-based PLpro ensemble

| PDB ID | State | PubChem CID | Resolution (Å) | PDB ID | State | PubChem CID | Resolution (Å) |
|---|---|---|---|---|---|---|---|
| 6W9C (C) | apo | - | 2.70 | 6XA9 (E) | apo | - | 2.90 |
| 6WUU (B) | holo | - | 2.79 | 7CJM | holo | 24941262 | 3.20 |
| 6WZU | apo | - | 1.79 | 7JRN (B) | holo | 24941262 | 2.48 |

**NSP12 - RNA-dependent RNA-polymerase (RdRp)**

Three ensembles were built for the RdRp protein, based on conformational changes in the catalytic binding site. The position of remdesivir in the crystallographic structure 7BV2 was taken as a reference for the targeted binding site (supplementary figure S5 A). At total, 7 crystallographic structures, out of 11 analysed, were included into the RdRp crystal ensemble (supplementary table S4). In addition, two new ensembles were produced by sampling conformations from molecular dynamics trajectories of RdRp in the APO state. At total 100,000 snapshots were analysed for each force field set of simulations. The amount of structures into the ensemble for crystals, charmm simulation and gromos simulations are 7, 14 and 13, respectively (supplementary figure S5 B-F). The scoring box center was set to 90.02, 90.17 and 103.52 (X, Y and Z, respectively) with dimensions 98x94x90 Å for crystal ensemble; 92.02, 90.17 and 103.52 (X, Y and Z, respectively) with dimensions 96x94x90 Å for charmm ensemble; and 92.02, 90.17 and 103.52 (X, Y and Z, respectively) with dimensions 96x94x90 Å for gromos ensemble. The box used for sampling was tuned to the binding site of each receptor conformation, based on the solvent accessible volume of the binding site.

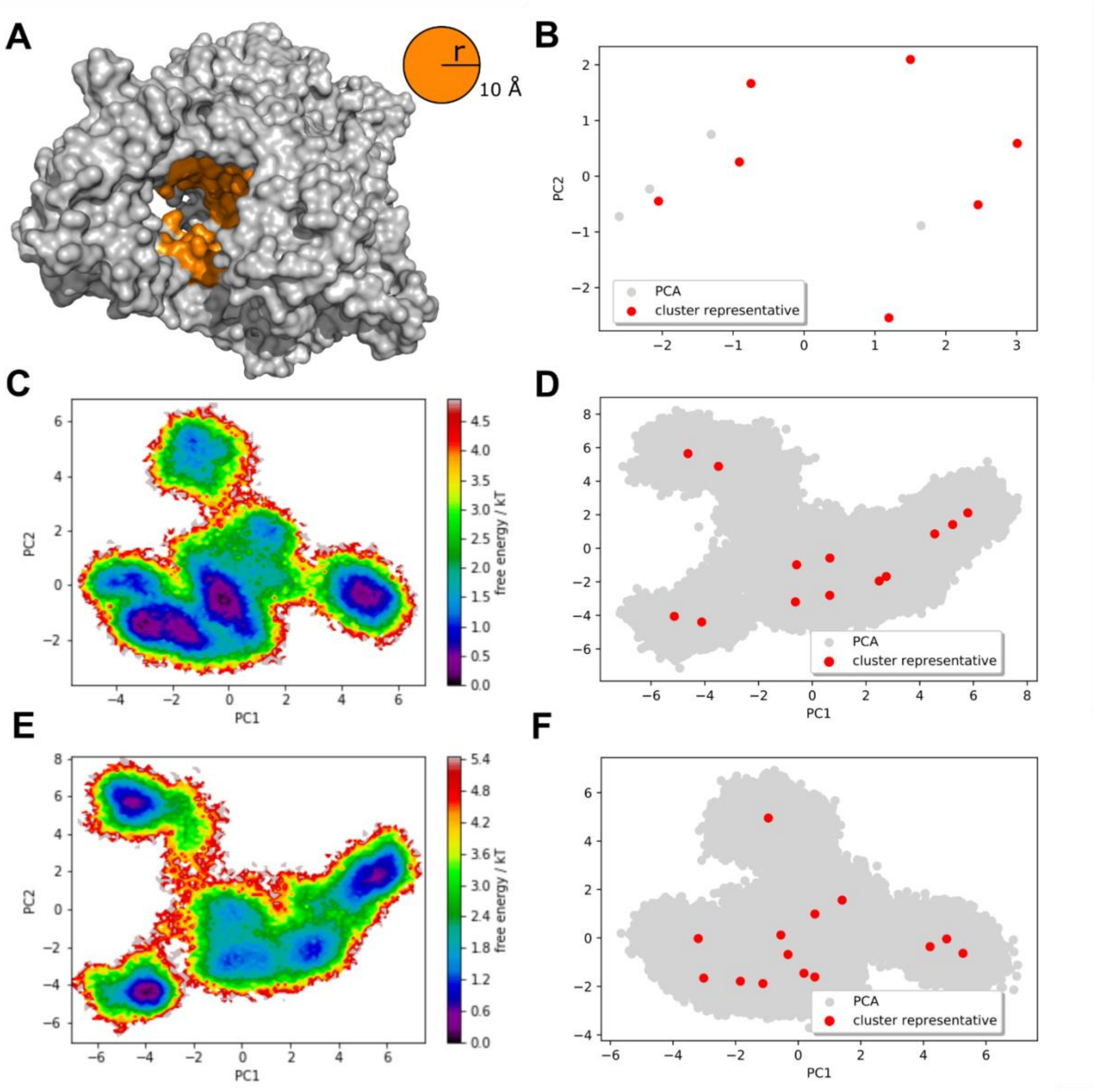


Supplementary Figure S6: Selection of representative conformations to create the RdRp ensembles. These ensembles take into account the flexibility of residues shown in orange (A). The conformational space for the unbound (i.e., apo) RdRp structure was sampled using data extracted from simulations using either available experimental structures (B), CHARMM36 force field (D) or GROMOS53a6 force field (F). In the case of MD, the initial input structure was modelled using the crystallographic data 7BTF as template. Approximately 100,000 conformations were extracted from MD trajectories. The K-means clustering algorithm from scikit-learn package (Pedregosa *et al.*, 2011)was employed to identify representative members for each cluster (red circles). The free energy of these conformations was estimated with PyEMMA (Scherer *et al.*, 2015)and plotted over the first two PCs (C, E) (i.e., darker colors indicate local minimas).

Supplementary table S4: Selected structures for the crystal-based RdRp ensemble

| PDB ID | State | PubChem CID | Resolution (Å) | PDB ID | State | PubChem CID | Resolution (Å) |
|---|---|---|---|---|---|---|---|
| 6M71 | apo | - | 2.90 | 7BW4 | apo | - | 3.70 |
| 6YYT | apo | - | 2.90 | 7BZF | apo | - | 3.26 |
| 7AAP | holo | 5271809 | 2.50 | 7D4F | holo | 365537 | 2.57 |
| 7BV1 | apo | - | 2.80 | | | | |

**Proof-of-concept validation with the Main Protease (Mpro)**

Although Mpro protein is, so far, one of the most explored SARS-CoV-2 targets in computational studies, there are still many open questions related to the design of appropriated and effective inhibitors. As pointed out by Kneller and collaborators (2020) (Kneller, Phillips, *et al.*, 2020), the malleability of the Mpro active-site cavity remains the greatest challenge in the development of effective inhibitors. While most of the available crystallographic data was collected at low temperature, thus limiting the flexibility of the binding pocket, x-ray data collected at room temperature was recently made available. This data demonstrates the ability of Mpro to substantially distort its shape and size in response to the presence of ligands (Kneller, Galanie, *et al.*, 2020). Such malleability allows this protein to fit accordingly to the diversity of physical-chemical features (i.e., chemical groups, size, charge distribution, etc) of ligands. Unfortunately, this ability is lost during the use of crystallographic structures as targets during virtual screenings using conventional docking approaches. In this work the conformational space of SARS-CoV-2 proteins was sampled from available crystallographic data, and from snapshots obtained through molecular dynamics simulations. Using a dataset of 36 small ligands it was possible to recover the near experimental conformation for the ligand (table S5), and show the ability of our ensemble docking protocol to explore the available conformational landscape within receptor ensembles (supplementary figure S7). In addition, tests with a set of 9 experimentally characterized SARS-CoV-2 Mpro inhibitors have shown the capability of DINC-COVID server to reproduce experimental data with significant strong correlation (e.g., r=0.88, r=0.8 and r=0.88 for crystals, charmm and gromos ensembles, respectively, see table S6). When compared with other two available docking web servers, DINC-COVID outperformed DockThor (Costa *et al.*, 2020) (r= 0.70), and showed similar performance to covid-19 (Kong *et al.*, 2020) (r=0.87). Nevertheless, the most significant difference was observed when antiviral compounds Leupeptin, Telaprevir, Narlaprevir and Boceprevir were tested. By the time the MPro crystal ensemble for DINC-COVID was build there were no experimental structures of Mpro bound to these compounds. Indeed, it was only recently that the crystallographic structure of these compounds in complex with SARS-CoV-2 MPro (6XCH, 6XQS, 6XQT and 6XQU) were solved at room temperature, thus allowing for the induced-fit process to occur without the constraints of crystallographic packing at very low temperatures. In this work, we show that by using Mpro conformations obtained from molecular dynamics simulations we were able to correlate our calculated binding energy with experimental binding affinity data for this challenging dataset (r=0.86). A comparison of our results with those of other docking servers is provided in supplementary table S7.

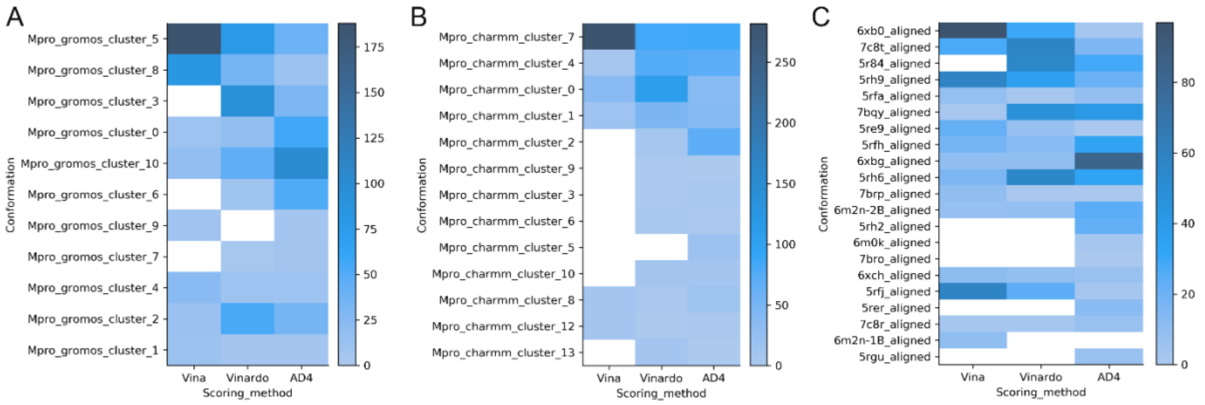


Supplementary figure S7: Distribution of receptor conformations included in the DINC-COVID outputs. Even when performing ensemble docking with a somewhat homogeneous dataset of small ligands, as the 36 drugs used in our validation experiment (see supplementary table S5), DINC-COVID explores a range of receptor conformations within the top scoring binding modes for each of the scoring functions used for rescoring. The heatmaps above show the number of times each receptor conformation is included in the top 15 best ranked binding modes produced by the ensemble docking of the 36 ligands using the GROMOS ensemble (A), the CHARMM ensemble (B), or the crystal ensemble (C).

**Supplementary table S5:** Self-docking results

| PDB ID | PubChem CID | DoFs[b] | LRMSDc (Å) | PDB ID | PubChem CID | DoFs[b] | LRMSDc (Å) |
|---|---|---|---|---|---|---|---|
| 5R7Y | 118569 | 4 | 1.91 | 5RG1 | 146019222 | 7 | 2.09 |
| 5R7Z | 405042899 | 3 | 1.49 | 5RGH | 125137283 | 2 | 2.43 |
| 5R80 | 89847 | 4 | 0.58 | 5RGI | 47172017 | 3 | 0.46 |
| 5R81 | 404847064 | 2 | 0.79 | 5RGK | 935612 | 4 | 3.59 |
| 5R82 | 24701445 | 2 | 2.15 | 5RGU | 146037571 | 5 | 1.31 |
| 5R83 | 674807 | 2 | 1.4 | 5RGV | 146037572 | 3 | 2.09 |
| 5R84 | 1072430 | 3 | 0.89 | 5RGW | 146037573 | 3 | 1.18 |
| 5RE4 | 20786326 | 1 | 0.61 | 5RGX | 146037574 | 3 | 0.51 |
| 5RE9 | 880785 | 3 | 2.57 | 5RGY | 146037575 | 4 | 0.91 |
| 5REB | 40476772 | 3 | 1.89 | 5RGZ | 146037576 | 3 | 0.63 |
| 5REH | 712045 | 4 | 2.79 | 5RH0 | 146037577 | 2 | 2.4 |
| 5REZ | 145998217 | 2 | 1.15 | 5RH1 | 146037578 | 3 | 1.29 |
| 5RF1 | 69696 | 2 | 1.45 | 5RH2 | 89468951 | 3 | 0.67 |
| 5RF2 | 6346752 | 2 | 1.03 | 5RH3 | 146037579 | 3 | 0.53 |
| 5RF3 | 344373 | 1 | 1.24 | 5RH8 | 146037583 | 5 | 1.03 |
| 5RF6 | 79838750 | 1 | 0.64 | 5RHD | 735904 | 2 | 0.49 |
| 5RF7 | 145998218 | 2 | 0.5 | 6M2N | 5281605 | 4 | 2.71 |
| 5RFE | 2815672 | 2 | 2.62 | 6W63 | 145998279 | 7 | 0.54 |
| | | | | **Mean** | | **3** | **1.4** |

This subset of 36 conformations was selected from a total of 156 crystal structures, after excluding covalently-bound ligands and apo (i.e., unbound) conformations. [b] DoFs, Degrees of Freedom (i.e., number of rotatable bonds). [c] LRMSD, Lower Root Mean Square Deviation of all heavy atoms of the predicted ligand, in relation to the crystal structure. Only the lowest value for each ligand is reported (top RMSD). Note that the goal of ensemble-docking is not to reproduce rigid crystal structures, but to find alternative lower energy binding modes. Therefore, although capable of sampling binding modes close to those observed in crystal structures, DIINC-COVID will often identify alternative binding modes with lower binding energy, taking advantage of the alternative receptor conformations available in the ensemble.

**Supplementary table S6:** Correlation between experimentally characterized inhibition values of Mpro drug-like inhibitors and docking-derived binding energies.

| | | | DINC-COVID (Crystals) | | | DINC-COVID (Charmm36) | | | DINC-COVID (Gromos53a6) | | | Covid-19 Server | DockThor server | |
|---|---|---|---|---|---|---|---|---|---|---|---|---|---|---|
| **Name** | **IC50** | **$\Delta G_{EXP}$[a]** | **Vina** | **Vinardo** | **AD4** | **Vina** | **Vinardo** | **AD4** | **Vina** | **Vinardo** | **AD4** | **TOP1** | **6LU7** | **6W63** |
| 11r | 0.18 | -9.23 | -8.92 | -10.8 | -15.29 | -8.44 | -8.82 | -13.94 | -7.95 | -8.88 | -13.09 | -7.40 | -9.159 | -9.471 |
| 13b | 0.67 | -8.45 | -8.51 | -9.82 | -14.2 | -8.11 | -9.14 | -12.46 | -7.92 | -8.12 | -12.65 | -7.40 | -8.698 | -8.788 |
| 11a | 0.053 | -9.96 | -8.86 | -9.66 | -12.62 | -8.21 | -8.58 | -10.27 | -8.08 | -8.18 | -10.66 | -8.10 | -7.976 | -8.771 |
| 11b | 0.04 | -10.1 | -9.08 | -9.98 | -12.03 | -8.29 | -8.25 | -10.46 | -8.09 | -8.5 | -9.89 | -8.10 | -8.245 | -8.627 |
| Carmofur | 1.82 | -7.86 | -6.68 | -7.18 | -6.08 | -6.48 | -6.28 | -5.49 | -6.1 | -5.83 | -4.71 | -6.20 | -7.306 | -7.09 |
| Disulfiram | 9.35 | -6.89 | -4.52 | -5.02 | -4.78 | -4.18 | -4.29 | -3.95 | -4.12 | -4.2 | -4.04 | -4.40 | -7.254 | -7.662 |
| Ebselen | 0.67 | -8.45 | -6.82 | -6.02 | -6.38 | -6.41 | -5.75 | -6.02 | -6.14 | -4.96 | -5.31 | -6.60 | -8.267 | -7.958 |
| PX-12 | 21.39 | -6.39 | -4.72 | -5.12 | -4.7 | -4.47 | -4.88 | -4.42 | -4.42 | -5.04 | -4.86 | -4.20 | -7.471 | -7.604 |
| Tideglusib | 1.55 | -7.95 | -8.43 | -8.28 | -9.38 | -8.68 | -7.12 | -7.43 | -7.46 | -6.31 | -6.87 | -8.00 | -7.944 | -8.287 |
| Correlation | | | **0.88** | 0.84 | 0.78 | 0.80 | 0.81 | 0.76 | **0.88** | 0.83 | 0.72 | 0.87 | 0.62 | 0.70 |

[a] $\Delta G_{EXP}$ was calculated using the equation $\Delta G_{EXP}$=RTln($k_i$). R is the gas constant, T is the temperature at 298 K, and ki is assumed to be equal to the experimental IC50 value (Zhang *et al.*, 2020; Jin, Du, *et al.*, 2020; Jin, Zhao, *et al.*, 2020; Dai *et al.*, 2020). The unit of energy is kcal $mol^{-1}$. The unit of IC50 is µM. The best Person correlation is highlighted in bold.

**Supplementary table S7:** Correlation between *in vitro* inhibition values of Mpro antiviral peptidomimetics and docking-derived binding energies.

| | | | DINC-COVID (Crystals) | | | DINC-COVID (Charmm36) | | | DINC-COVID (Gromos53a6) | | | Covid-19 Server | DockThor server | |
|---|---|---|---|---|---|---|---|---|---|---|---|---|---|---|
| **Name** | **IC50** | **$\Delta G_{EXP}$[a]** | **Vina** | **Vinardo** | **AD4** | **Vina** | **Vinardo** | **AD4** | **Vina** | **Vinardo** | **AD4** | **TOP1** | **6LU7** | **6W63** |
| Telaprevir | 18 | -6.5 | -9.08 | -9.64 | -14.6 | -7.98 | -8.72 | -13.1 | -8.12 | -8.8 | -12.68 | -8.40 | -8.998 | -8.904 |
| Leupeptin | 92 | -5.5 | -6.94 | -7.96 | -9.79 | -6.57 | -7.37 | -8.77 | -6.32 | -6.72 | -8.6 | -6.10 | -7.152 | -8.459 |
| Boceprevir | 3.1 | -7.5 | -8.33 | -8.12 | -12.33 | -7.95 | -7.45 | -10.55 | -7.6 | -7.09 | -10.41 | -7.20 | -7.804 | -8.767 |
| Narlaprevir | 5.1 | -7.2 | -8.25 | -9.35 | -16.81 | -7.91 | -7.27 | -15.03 | -7.22 | -7.77 | -12.42 | -6.80 | -9.161 | -7.873 |
| Correlation | | | 0.60 | 0.24 | 0.61 | **0.86** | -0.14 | 0.54 | 0.58 | 0.18 | 0.53 | 0.32 | 0.48 | -0.12 |

[a] $\Delta G_{EXP}$ was calculated using the equation $\Delta G_{EXP}$=RTln($k_i$). R is the gas constant, T is the temperature at 298 K, and *ki* is assumed to be equal to the experimental IC50 value (Kneller, Galanie, *et al.*, 2020). The unit of energy is kcal $mol^{-1}$. The unit of IC50 is µM. The best Person correlation is highlighted in bold.

**References:**


Best,R.B. *et al.* (2012) Optimization of the additive CHARMM all-atom protein force field targeting improved sampling of the backbone ϕ, ψ and side-chain χ1 and χ2 Dihedral Angles. *J. Chem. Theory Comput.*

Costa,L.S.C. *et al.* (2020) Drug Design and Repurposing with DockThor-VS Web Server: Virtual Screening focusing on SARS-CoV-2 Therapeutic Targets and their Non-Synonym Variants. *Res. Sq.*

Dai,W. *et al.* (2020) Structure-based design of antiviral drug candidates targeting the SARS-CoV-2 main protease. *Science (80-. ).*

Dolinsky,T.J. *et al.* (2004) PDB2PQR: An automated pipeline for the setup of Poisson-Boltzmann electrostatics calculations. *Nucleic Acids Res.*, **32**.

Jin,Z., Zhao,Y., *et al.* (2020) Structural basis for the inhibition of SARS-CoV-2 main protease by antineoplastic drug carmofur. *Nat. Struct. Mol. Biol.*

Jin,Z., Du,X., *et al.* (2020) Structure of Mpro from SARS-CoV-2 and discovery of its inhibitors. *Nature*.

Kneller,D.W., Galanie,S., *et al.* (2020) Malleability of the SARS-CoV-2 3CL Mpro Active-Site Cavity Facilitates Binding of Clinical Antivirals. *Structure*.

Kneller,D.W., Phillips,G., *et al.* (2020) Structural plasticity of SARS-CoV-2 3CL Mpro active site cavity revealed by room temperature X-ray crystallography. *Nat. Commun.*

Kong,R. *et al.* (2020) COVID-19 Docking Server: An interactive server for docking small molecules, peptides and antibodies against potential targets of COVID-19. *arXiv*.

Krissinel,E. and Henrick,K. (2007) Inference of Macromolecular Assemblies from Crystalline State. *J. Mol. Biol.*, **372**, 774–797.

Lopes,P.E.M. *et al.* (2015) Current status of protein force fields for molecular dynamics simulations. *Methods Mol. Biol.*

McGibbon,R.T. *et al.* (2015) MDTraj: A Modern Open Library for the Analysis of Molecular Dynamics Trajectories. *Biophys. J.*

Menéndez,C.A. *et al.* (2020) Molecular characterization of ebselen binding activity to SARS-CoV-2 main protease. *Sci. Adv.*

Morris,G.M. *et al.* (2009) AutoDock4 and AutoDockTools4: Automated docking with selective receptor flexibility. *J. Comput. Chem.*, **30**, 2785–2791.

Olsson,M.H.M. *et al.* (2011) PROPKA3: Consistent Treatment of Internal and Surface Residues in Empirical p<italic>K</italic>$_a$ Predictions BT - Journal of Chemical Theory and Computation. *J. Chem. Theory Comput.*

Oostenbrink,C. *et al.* (2004) A biomolecular force field based on the free enthalpy of hydration and solvation: The GROMOS force-field parameter sets 53A5 and 53A6. *J. Comput. Chem.*

Pedregosa,F. *et al.* (2011) Scikit-learn: Machine learning in Python. *J. Mach. Learn. Res.*

Quiroga,R. and Villarreal,M.A. (2016) Vinardo: A scoring function based on autodock vina improves scoring, docking, and virtual screening. *PLoS One*.

Scherer,M.K. *et al.* (2015) PyEMMA 2: A Software Package for Estimation, Validation, and Analysis of Markov Models. *J. Chem. Theory Comput.*

Trott,O. and Olson,A.J. (2009) AutoDock Vina: Improving the speed and accuracy of docking with a new scoring function, efficient optimization, and multithreading. *J. Comput. Chem.*

Villavicencio,B. *et al.* (2018) All-Hydrocarbon Staples and Their Effect over Peptide Conformation under Different Force Fields. *J. Chem. Inf. Model.*

Zhang,L. *et al.* (2020) Crystal structure of SARS-CoV-2 main protease provides a basis for design of improved α-ketoamide inhibitors. *Science (80-. ).*, **368**, 409 LP – 412.
